# Synovial Fluid Mitochondrial DNA Concentration Reflects the Degree of Cartilage Damage After Naturally Occurring Articular Injury

**DOI:** 10.1101/2021.08.26.457571

**Authors:** Lindsay A. Seewald, Isabella G. Sabino, Kaylee L. Montney, Michelle L. Delco

## Abstract

Posttraumatic osteoarthritis (PTOA) is a debilitating sequela to joint injury with no current therapeutics that can slow its progression. Early intervention, prior to the development of degenerative joint changes, has the potential for greater therapeutic success but requires early detection of joint injury. In other tissue types, trauma is associated with the extracellular release of mitochondrial DNA (mtDNA), which serves as a mitochondria-specific Damage Associated Molecular Pattern (mDAMP) to perpetuate inflammation. We demonstrated that chondrocytes release mtDNA following cellular stress and that mtDNA is increased in equine synovial fluid following experimental and naturally occurring mechanical injury to the joint surface. Moreover, we found a strong correlation between the degree of cartilage damage and mtDNA concentration. Finally, impact-induced mtDNA release was mitigated by mitoprotective treatment. These data suggest synovial fluid mtDNA may represent a sensitive marker of early articular injury, prior to the onset of changes on standard diagnostic imaging modalities.

**One Sentence Summary:** Synovial fluid mitochondrial DNA increases after articular injury, is associated with the degree of cartilage damage, and is mitigated by mitoprotective treatment.

## Introduction

Post-traumatic osteoarthritis (PTOA) affects approximately 5.6 million individuals in the United States *(1)*, yet current therapies are unable to modify the progression of disease *(2, 3)*. Mounting evidence suggests that targeting very early pathomechanisms after articular injury may be the key to preventing osteoarthritis (OA) *(4)*; however, clinical signs of joint pain and loss of mobility often appear years or decades after the inciting trauma, making early detection and intervention difficult or impossible *(5)*. Furthermore, current diagnostics, including radiographs and MRI, often are not sensitive enough to detect disease prior to the onset of structural changes in the joint *(6)*. Therefore, a thorough understanding of the response of cartilage to mechanical injury and the events that perpetuate damage signals throughout the joint is essential to the development of more sensitive diagnostic tests and effective therapies for PTOA *(6, 7)*.

Recent work has demonstrated that mitochondrial dysfunction occurs as an acute response of chondrocytes to mechanical injury, resulting in several processes that lead to the development of PTOA, including the production of proinflammatory cytokines, chemokines, and proteases *(8)*; a reduction in proteoglycan and collagen synthesis *(9, 10)*; glycosaminoglycan (GAG) release *(9, 11)*; and cartilage calcification *(10)*. Thus, mitochondrial dysfunction represents a key event that could serve as an indicator of cartilage damage in the peracute time frame after injury. In addition, preserving mitochondrial structure and function, termed mitoprotection, may represent a valuable new strategy for preventing PTOA progression. Szeto-Schiller (SS) peptides are a class of mitoprotective agents that repair cristae structure and restore mitochondrial function, thereby improving cellular bioenergetics and preventing cell death *(12–14)*. In an *ex vivo* cartilage impact model of PTOA, treatment with the mitoprotective peptide, SS-31 (elamipretide), shortly after injury prevented chondrocyte death and cartilage matrix degradation *(11)*. The clinical application of mitoprotective agents such as SS-31 to treat acute cartilage injury and prevent the onset of PTOA would require a practical and sensitive diagnostic test to identify patients experiencing mitochondrial dysfunction and to monitor response to mitoprotective therapy. Unfortunately, no such disease markers have yet been identified.

One class of molecules, mitochondria-specific Damage Associated Molecular Patterns (mDAMPS), may prove to be useful indicators of early joint injury and mitochondrial dysfunction. mDAMPs are molecules such as mitochondrial DNA (mtDNA), cardiolipin, N-formyl peptide, and cytochrome c that are contained exclusively within the mitochondria in a healthy cell *(15)*. When released into the cytosol or the extracellular space, mDAMPs drive pro-inflammatory responses via a variety of pathways *(16)*. One particular mDAMP, mtDNA, has been validated as a biomarker of cellular stress in several tissue types and disease states *(17–19)*. Of particular relevance to orthopedic tissues, synovial fluid mtDNA was positively correlated with the degree of inflammation in rheumatoid arthritis patients *(20, 21)* and elevated serum mtDNA has been associated with femoral fractures *(22)*. As extracellular mtDNA has been associated with traumatic injury to cells *(17, 22, 23)*, synovial fluid mtDNA may be a promising candidate for early detection of orthopedic conditions precipitated by mechanical injury, such as PTOA.

Several potential mechanisms for extracellular mtDNA release have been proposed, including opening of the mitochondrial permeability transition pore (MPTP) *(24, 25)*, altered mitophagy *(26)*, and release of exosomes *(27, 28)*. These mechanisms occur concurrently with reactive oxygen species (ROS) formation and mitochondrial dysfunction *(26, 29, 30)*, suggesting that extracellular mtDNA may be an indicator of mitochondrial dysfunction within cells. In the context of PTOA, synovial fluid extracellular mtDNA may provide a practical method for early diagnosis of cartilage injury and facilitate monitoring of response to mitoprotective treatment. Here, we use a robust series of translational models, from *in vitro* cell culture to clinical cases of equine intra-articular fracture, to investigate extracellular mtDNA release and the effects of mitoprotective therapy following acute cartilage injury.

## Results

### Chondrocytes release mtDNA in response to an inflammatory stimulus

To evaluate whether chondrocytes release mtDNA following cellular stress, we used an established *in vitro* cell culture model of osteoarthritis *(31)*. Equine chondrocytes were harvested from the femoropatellar joints of healthy, young adult horses (n = 4). *In vivo*, articular chondrocytes are adapted to low oxygen tension and low glucose concentrations compared to most tissues, and these conditions influence chondrocyte mitochondrial function and metabolic state *(32–34)*. Therefore, to simulate a physiologic environment, chondrocytes were cultured at 5% oxygen and 5 μM glucose. To model osteoarthritis, we stimulated second passage chondrocytes with a sublethal dose of the inflammatory mediator interleukin 1 beta (IL-1β, 1 ng/μL). Total cell counts 12 and 24 hours after addition of stimulant media were not different between unstimulated and stimulated cells (Fig. 1A), confirming that IL-1β did not result in significant chondrocyte death.

**Fig. 1.**
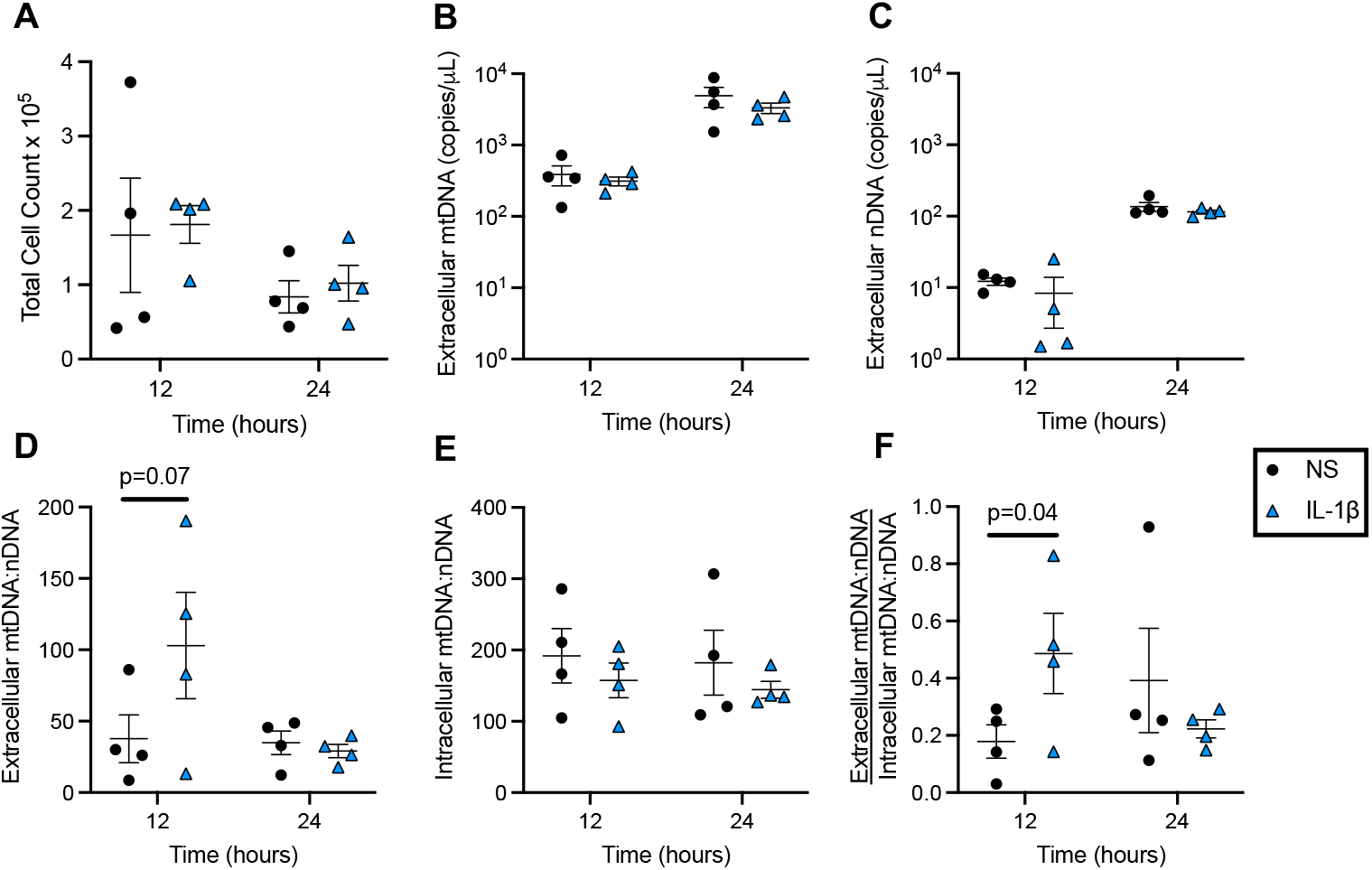
Cultured chondrocytes selectively release mtDNA in response to an inflammatory stimulus. (**A**) Chondrocyte number assessed at 12 and 24 hours after the start of IL-1β stimulation. (**B-C**) Extracellular mtDNA and nDNA concentration in chondrocyte-conditioned media. (**D**) Extracellular mtDNA concentration normalized to extracellular nDNA concentration (extracellular mtDNA:nDNA ratio). (**E**) Intracellular mtDNA concentration normalized to intracellular nDNA concentration (intracellular mtDNA:nDNA ratio). (**F**) Extracellular mtDNA:nDNA ratio normalized to intracellular mtDNA:nDNA ratio. All data are means ± SEM, n = 4, statistics by students’ (paired) t test. Line connects groups different with p-values listed.

At 12 and 24 hours after stimulation, extracellular DNA content in chondrocyte-conditioned media and intracellular DNA content in cultured chondrocytes were assessed using quantitative PCR (qPCR). Extracellular nuclear DNA concentration (nDNA) was quantified in addition to mtDNA in order to differentiate between mtDNA released secondary to cell death, versus mtDNA released via active mechanisms from live, stressed chondrocytes; nDNA is released during apoptosis and necrosis *(35)*, so nDNA is considered an indicator of cell death, while mtDNA release can occur passively during cell death or actively by viable cells *(35)*. Thus, quantifying extracellular nDNA, mtDNA, and the mtDNA:nDNA ratio allowed us to differentiate between active and passive mechanisms of mtDNA release. We found that neither extracellular mtDNA (Fig. 1B), nDNA (Fig. 1C), nor extracellular mtDNA:nDNA ratio (Fig. 1D) were significantly different between IL-1β-stimulated wells and controls. However, since studies suggest that the intracellular mtDNA:nDNA ratio can vary due to a variety of factors including aging, metabolic state, or oxidative stress/mitochondrial dysfunction *(36)*, we quantified intracellular mtDNA concentrations. We found that the intracellular mtDNA:nDNA ratio did not vary between groups or time points (Fig. 1E) but did vary between individuals and chondrocyte passage (fig. S1). To account for this biologic variability, we normalized extracellular mtDNA:nDNA ratio to intracellular mtDNA:nDNA ratio for each individual. This normalized extracellular mtDNA:nDNA ratio was significantly elevated in IL-1β-stimulated chondrocytes compared to controls at 12 hours after stimulation (Fig. 1E, p = 0.04). This relative increase in extracellular mtDNA compared to intracellular mtDNA suggests that live chondrocytes selectively release mtDNA in response to inflammatory stress.

### Mitoprotection mitigates mtDNA release following mechanical stress ex vivo

The main factor that contributes to the development of PTOA is mechanical injury to the articular surface, which can occur either at the time of trauma or as a result of altered joint loading following injury *(6, 37)*. Therefore, we investigated whether injurious mechanical loading of cartilage would result in mtDNA release from chondrocytes. Previously, we demonstrated that mechanical impact injury to cartilage explants induces mitochondrial dysfunction, chondrocyte apoptosis, and cartilage matrix degeneration *(11, 38)*. Using a similar study design, we applied a single rapid compression (impact) injury to bovine cartilage explants and quantified extracellular mtDNA and nDNA from cartilage conditioned media for several days following injury (Fig. 2A). In addition, we treated one group of unimpacted and one group of impacted explants with the mitoprotective peptide SS-31 at either 1 hour or 12 hours after injury to investigate the relationship between mitochondrial dysfunction and extracellular DNA release.

**Fig. 2.**
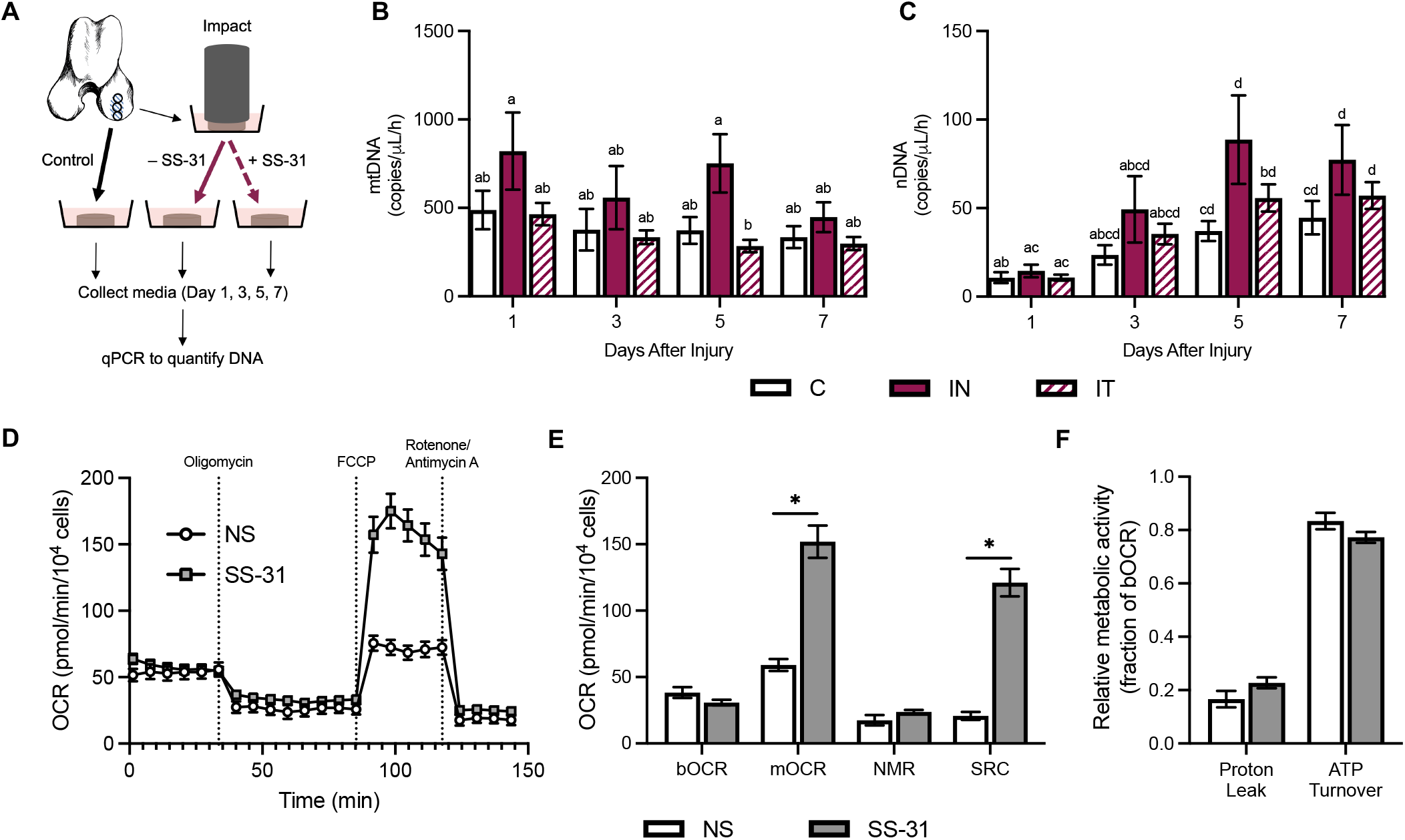
Mitoprotection mitigates mtDNA release from chondrocytes following mechanical injury *ex vivo*. (**A**) Cartilage explants were harvested from the medial femoral condyle of neonatal bovids and injured by delivering a single, rapid mechanical overload (impact). Impacted explants were cultured for 7 days without treatment (IN, n = 9) or with mitoprotective (SS-31) treatment (IT, n = 16). Control (C, n = 6) explants received neither impact nor treatment. Cartilage conditioned media was collected serially for qPCR quantification of extracellular mtDNA and nDNA. (**B-C**) Conditioned media mtDNA and nDNA concentration for control (C), injured untreated (IN) and injured treated (IT) explants. (**D**) Chondrocyte oxygen consumption rate (OCR) during mitochondrial stress test with (SS-31) and without (NS) mitoprotective treatment (n = 7). (**E**) Basal OCR (bOCR), maximal OCR (mOCR), non-mitochondrial respiration (NMR), and spare respiratory capacity (SRC) for chondrocytes with and without treatment. (n = 7). (**F**) Proton leak and ATP turnover (as a fraction of bOCR) for chondrocytes with and without treatment (n = 7). All data are means ± SEM. (**B-C)** Statistics by linear mixed effects model. Groups not sharing a letter are significantly different (p < 0.05). (**E-F**) Statistics by students’ t test. *P < 0.05 between groups.

Neither extracellular mtDNA, nDNA, nor mtDNA:nDNA ratio were significantly different between impacted (IN, n = 9) and unimpacted (C, n = 6) groups at the time points measured (Fig. 2B). However, within each group cell death increased over the course of the study (Fig. 2C, p < 0.05), suggesting that unimpacted control samples experienced cellular stress in this *ex vivo* tissue culture model. This ongoing cell death likely makes detection of active mtDNA release difficult and may explain why we did not see differences in mtDNA:nDNA ratio between groups in this model. Within each treatment group, the extracellular mtDNA:nDNA ratio was significantly higher on day one than at later time points (fig. S2, p < 0.05), which likely reflects either a greater proportion of active release of mtDNA in the acute phase after injury, or a decrease in intracellular mtDNA content over the course of the experiment due to mitochondrial dysfunction-induced mitophagy *(30)*.

We next investigated the effects of mitoprotection on cartilage explants. While extracellular mtDNA from impacted cartilage (IN, n = 9) were not significantly different from controls (C, n = 6), SS-31 treatment of impacted explants (IT, n = 16) produced a significant decrease in mtDNA concentration compared to the untreated explants (IN) five days after injury (Fig. 2B, p < 0.05). Taken together, these findings suggest that mitochondrial dysfunction occurs concurrently with mtDNA release. To further characterize the effects of SS-31 on mitochondrial function, we conducted microscale respirometry on cultured chondrocytes with and without SS-31 treatment. Chondrocytes treated with SS-31 displayed a higher maximum oxygen consumption rate (mOCR) and spare respiratory capacity (SRC) than untreated chondrocytes (Fig. 2D-E, p < 0.05). There was no difference in the relative contributions of proton leak and ATP turnover to basal oxygen consumption rate (bOCR) (Fig. 2F, p < 0.05). mOCR and SRC are measures of the cell’s ability to increase ATP production to meet an increase in cellular energy demand *(39)*. In the face of cellular stressors including mechanical injury, the energy requirements of chondrocytes increase *(40)*; thus, the ability of SS-31 to increase SRC and improve chondrocytes’ ability to respond to cellular injury likely explains the current findings, as well as our previous findings that mitoprotection prevents impact-induced chondrocyte mitochondrial depolarization, apoptosis, and cartilage matrix degradation *(11, 41)*.

Finally, we investigated whether the timing of mitoprotective treatment affected mtDNA release. SS-31 treatment to 12 hours after injury (n = 6) was as protective against mtDNA release as treatment 1 hour after injury (n = 10) (fig. S3, p < 0.05). Similarly, explants treated at 1 and 12 hours after injury experienced similar amounts of cell death (fig. S3). In a previous study, we found no difference in cartilage degeneration or chondrocyte death when treatment was delayed to 12 hours after injury *(11)*. Our mtDNA and nDNA data support these previous findings and suggest that mitoprotection with SS-31 may have a therapeutic window of at least 12 hours following articular injury to prevent mitochondrial dysfunction, chondrocyte death, and subsequent cartilage degeneration in this *ex vivo* model.

### Mitoprotection mitigates mtDNA release following impact-injury in vivo

After measuring mtDNA release following mechanical stress in an ex vivo cartilage explant model, we investigated whether a similar injury would produce a change in synovial fluid mtDNA concentration *in vivo*. Using a previously validated equine model of early PTOA *(42)*, focal impact injuries of defined magnitude were delivered to the articular surface of the medial trochlear ridge of adult horses (n = 6) under arthroscopic guidance (Fig. 3A-C, movie S1). The location of this experimental injury is analogous to the most common site of lesions on the human talus leading to talocrural joint OA *(37, 43)* (Fig. 3B). In this study, experimentally applied articular impacts produced an elevation in mtDNA 7 days after injury compared to baseline concentrations (Fig. 3D), indicating that changes in synovial fluid mtDNA concentrations are detectable following cartilage injury *in vivo*. In addition, this model produced considerable cell death, as indicated by elevated nDNA concentrations, and decreased mtDNA:nDNA ratios at all time points after injury (Fig. 3E-F).

**Fig. 3.**
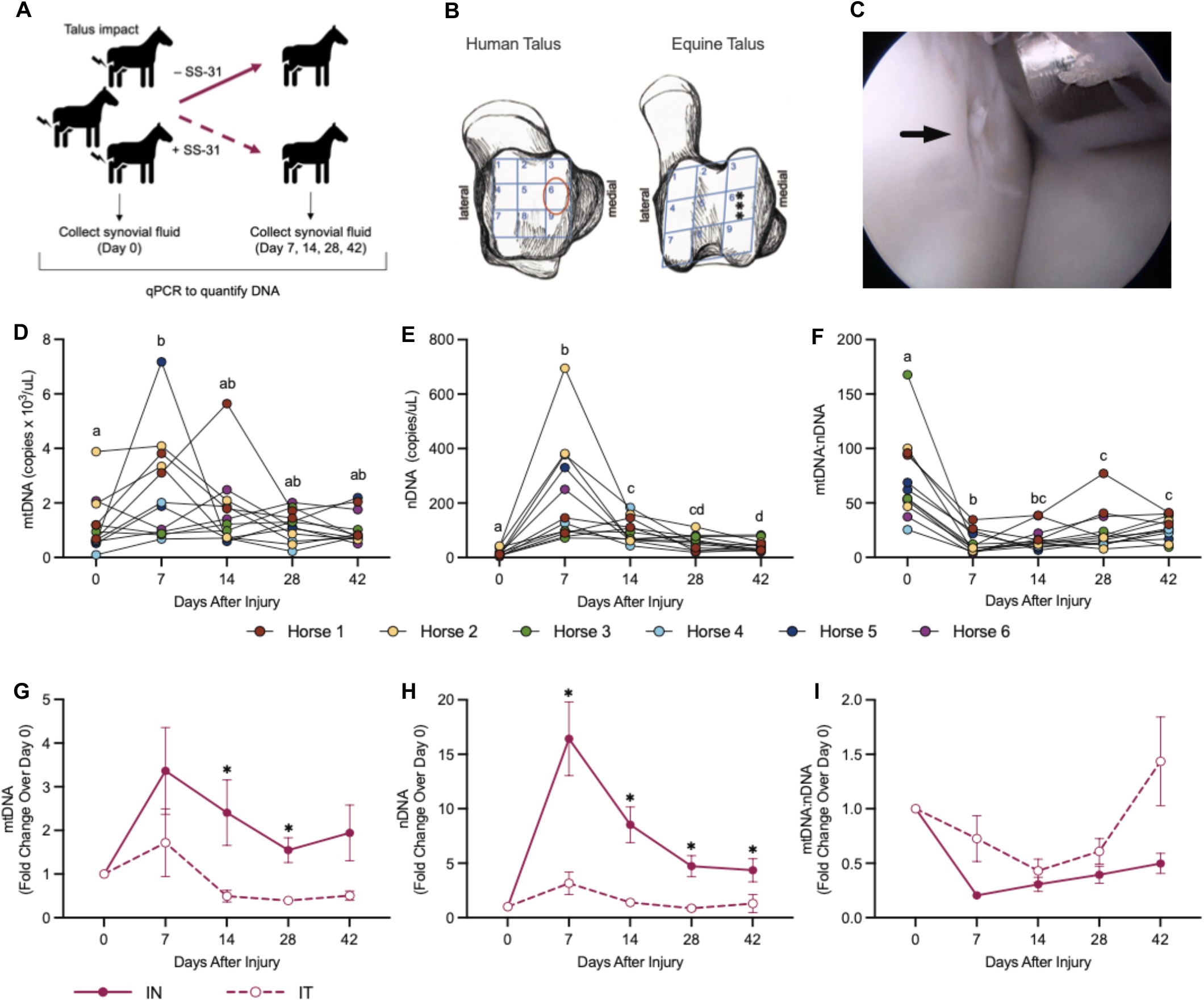
Mitoprotection mitigates mtDNA release following mechanical articular cartilage injury *in vivo*. (**A**) Adult horses were placed under general anesthesia and three focal cartilage injuries were delivered to the talus in each talocrural joint. One group was treated with intra-articular injection of the mitoprotective peptide SS-31 one hour after injury. Synovial fluid was collected serially after injury and compared to pre-injury synovial fluid samples. (**B**) Schematic comparing the anatomy of the human and equine talus. Asterisks indicate sites of cartilage impact. Red circle identifies analogous region on the human talus *(from Delco, et al*., *AJSM 2020; permission pending)*. (**C**) Arthroscopic image of the impacting device positioned on the medial trochlear ridge of the talus after impact. Arrow indicates cartilage defect caused by impact. (**D-F)** Synovial fluid mtDNA, nDNA, and mtDNA:nDNA ratio in injured, untreated joints (IN, n = 12). Time points not sharing a letter are significantly different (p < 0.05), statistics by linear mixed effects model. (**G-I)** Fold change over baseline (day 0, preinjury) of synovial fluid mtDNA, nDNA, and mtDNA:nDNA ratio in untreated (IN, n = 12) versus treated (IT, n = 10) joints. Data are means ± SEM, statistics by linear mixed effects model. *P<0.05 compared to IT within time points.

To evaluate the effects of mitoprotection *in vivo*, a second, randomly assigned group of horses (n=5) were delivered the same impact injury, then treated with a single intra-articular dose of SS-31 one hour after injury. Mitoprotective therapy significantly reduced concentrations of synovial fluid mtDNA compared to untreated joints at 14 and 28 days after injury (Fig. 3G). This finding suggests that injury-associated elevations in mtDNA are prevented by early mitoprotective intervention, and synovial fluid DNA may be a useful measure of response to mitoprotection *in vivo*. Further, cell death was significantly reduced in SS-31 treated joints at all time points during the study (Fig. 3H). No differences in mtDNA:nDNA ratio were observed between treated and untreated joints (Fig. 3I).

### Synovial fluid mtDNA correlates with severity of cartilage damage in naturally occurring joint injury

Next, we investigated synovial fluid mtDNA in cases of naturally occurring articular injury. For our study group, we chose equine patients presenting for intra-articular osteochondral carpal fragment removal, since these injuries are common in racehorses *(44)*, are traumatic in origin resulting from either repetitive loading or a single acute overload, and frequently lead to development of PTOA *(45)*. Synovial fluid samples were analyzed from a total of 19 horses with intra-articular osteochondral fragments. Patients were predominantly male (84%), 3–5-year-old (79%) Thoroughbred (79%) racehorses (74%) presenting for chip fractures involving either the radiocarpal joint (n = 6, 32%) or the middle carpal joint (n = 13, 68%). For comparison, synovial fluid was analyzed from the radiocarpal or middle carpal joints of clinically normal horses (n = 7 horses, n = 12 joints). There was no significant difference in age or sex distribution between the normal and fracture groups; however, differences in breed did exist between the groups, with the fracture group consisting of a higher proportion of thoroughbreds (p = 0.0002) than the control group. This difference is attributed to the difficulty in obtaining samples from healthy joints of elite thoroughbred athletes. See table S1 for detailed patient and sample information.

In agreement with our experimental injury model, synovial fluid mtDNA and nDNA concentrations were elevated in injured joints compared to normal joints (Fig. 4A and 4B). No significant differences existed between patient age, breed, sex, or side of injury and mtDNA concentration, nDNA concentration, or mtDNA:nDNA ratio (table S2). Among injured joints, mtDNA but not nDNA concentrations were higher in middle carpal joint fractures compared to radiocarpal joint fractures (Fig. 4C and 4D). There were no significant differences in the mtDNA:nDNA ratio between injured and normal joints or between middle carpal and radiocarpal fractures (fig. S4). In agreement with our *in vivo* articular injury model, these findings suggest that absolute mtDNA concentration in synovial fluid may be a more clinically useful measure of joint injury than mtDNA:nDNA ratio.

**Fig. 4.**
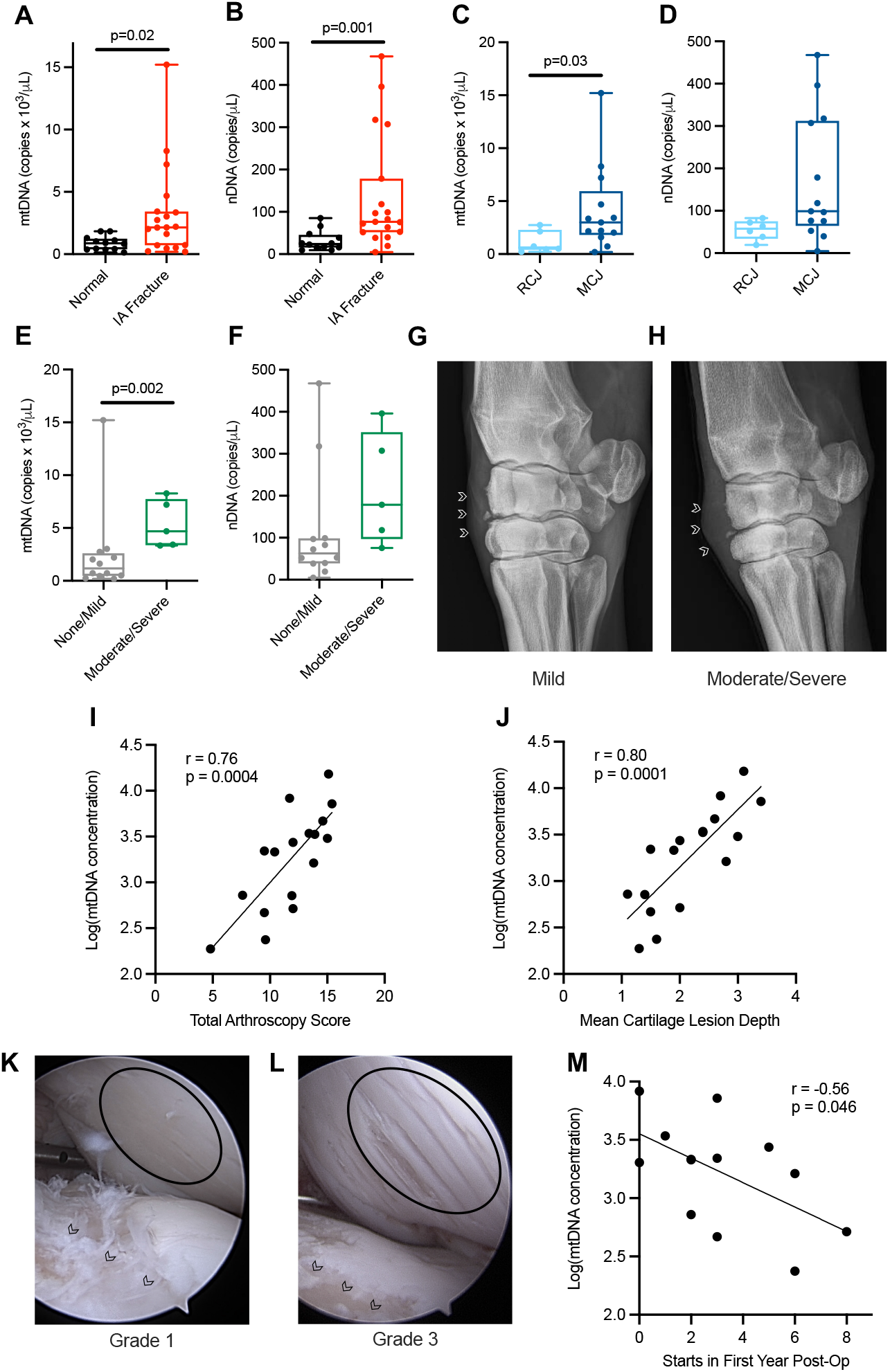
Synovial fluid mtDNA concentration correlates with the severity of cartilage injury. (**A-B)** Synovial fluid mtDNA and nDNA concentration in joints with intra-articular (IA) carpal fractures compared to normal joints. (**C-D**) Comparison of mtDNA and nDNA concentration between radiocarpal joint (RCJ) and middle carpal joint (MCJ) fractures. (**E-F**) Synovial fluid mtDNA and nDNA concentrations in joints with intra-articular carpal fractures and no or mild soft tissue swelling/joint effusion compared to those with fractures and moderate to severe soft tissue swelling/joint effusion. (**G-H**) Representative radiographs of joints with mild and moderate to severe soft tissue swelling/joint effusion. (**I**) Correlation of total arthroscopy score with mtDNA concentration in joints with IA fractures. (**J**) Correlation of mean cartilage lesion depth with mtDNA concentration in joints with IA fractures. (**K-L**) Representative arthroscopic images of joints with grade 1 and grade 3 cartilage lesion depth. Circle indicates area of cartilage damage on articulating surface. Arrow heads indicate fracture bed. (**M**) Correlation of number of race starts in the first year after surgery with mtDNA concentration. (**A-F**) Statistics by students’ t test. Line connects groups that are significantly different with p-values listed. (**I, J, M**) Statistics by Pearson correlation analysis.

Standard radiographic views of injured joints were analyzed for several criteria associated with degenerative joint disease *(46–50)*, including osteophyte formation, subchondral bone sclerosis, subchondral bone lysis, joint space narrowing, fracture severity, and soft tissue swelling/joint effusion (table S3). Radiographic studies were available for 17 out of 19 of the injured joints. The mean total radiograph score for joints with intra-articular fractures was 5.1 (95% CI, 3.4-6.8) out of a maximum score of 15, indicating that the joints in this study displayed little radiographic evidence of joint disease. No significant correlations between any of the scoring categories and mtDNA concentration, nDNA concentration, or mtDNA:nDNA ratio were found, except for moderate negative correlations between mtDNA:nDNA ratio and fracture severity (r = -0.61, p = 0.008) and mtDNA:nDNA ratio and total radiograph score (r = -0.49, p = 0.046) (table S4). However, we found that joints with moderate to severe soft tissue swelling had significantly higher mtDNA but not nDNA or mtDNA:nDNA ratio, than joints with no or mild soft tissue swelling (Fig. 4E-H, fig. S4). Altogether, these data support the hypothesis that mtDNA concentration may be a clinically useful indicator of early joint disease, prior to the onset of radiographically detectable changes associated with OA.

To investigate the relationship between extracellular mtDNA and early degenerative change to soft tissues within the joint including cartilage and synovial membrane, arthroscopic video footage was scored for severity of cartilage lesion depth, cartilage lesion size, active synovitis, chronic changes to the synovium, and subchondral bone pathology (table S5). Video footage was available for 17 out of the 19 injured joints. A total arthroscopy score, calculated as the sum of the five sub-scores, showed a strong positive correlation (r = 0.76, p = 0.0004) with synovial fluid mtDNA and nDNA concentration (Fig. 4I, fig S5). In addition, we evaluated the correlation of each sub-score with mtDNA concentration. Cartilage lesion depth was scored at four sites: immediately surrounding the fracture site, on the same bone but more distant to the fracture site, on the opposing articular surface to the fracture, and in remote locations within the same joint, using a modified ICRS scoring system *(51)* (table S5, fig. S6). These four scores were averaged to obtain a mean cartilage lesion depth score for each joint, which showed the strongest correlation (r = 0.80, P = 0.0001) with mtDNA concentration of any of the scoring criteria (Fig. 4I-L). This finding suggests that for our patient cohort, synovial fluid mtDNA reflects the degree of cartilage damage within the injured joint. Several of the other scoring categories showed moderate correlations with either mtDNA or nDNA (Table 1, table S6). Interestingly, subchondral bone pathology had no correlation with either mtDNA or nDNA (Table 1), which, in agreement with our radiographic findings, may indicate that mtDNA release is associated with the soft tissues of the joint rather than bone. Finally, we measured post-operative performance in patients that were Thoroughbred racehorses (n = 14) by obtaining race records from a public database. Race records were available for 13/14 patients, and all (13/13) returned to racing after surgery. Moreover, synovial fluid mtDNA concentration showed a moderate negative correlation (r = -0.56, p = 0.046) with the number of race starts in the first year after surgery (Fig. 4M), suggesting that patients with lower mtDNA at the time of surgery may have improved post-operative outcomes.

**Table 1.** Arthroscopic scoring correlation data. Values with R > 0.7 and P < 0.001 are bolded. Statistics by Pearson’s correlation analysis.

|  | mtDNA |  | nDNA |  |
| --- | --- | --- | --- | --- |
|  | log(concentration) |  | log(concentration) |  |
|  | R | P | R | P |
| Cartilage Lesion Depth | **0.80** | **<0.001** | **0.75** | **<0.001** |
| Cartilage Lesion Size | 0.59 | 0.012 | 0.69 | <0.001 |
| Subchondral Bone | 0.30 | 0.237 | 0.19 | 0.456 |
| Active Synovitis | 0.64 | 0.006 | 0.61 | 0.010 |
| Chronic Synovial Change | 0.46 | 0.062 | 0.68 | 0.003 |
| Total | **0.76** | **<0.001** | **0.79** | **<0.001** |

## Discussion

This study aimed to investigate mtDNA release from articular chondrocytes following mechanical injury and to evaluate the effects of mitoprotective treatment on extracellular mtDNA concentrations. Our data indicate that mtDNA is released from chondrocytes in response to cellular stress and that mitoprotective therapy can prevent this release. While extracellular mtDNA has previously been investigated as a potential biomarker in several other tissue types and disease states *(17, 18, 21–23, 27, 52, 53)*, this is the first evidence that chondrocytes release mtDNA and that increases in mtDNA can be detected in synovial fluid following articular injury. These findings are important because they suggest mtDNA quantification may represent a valuable diagnostic tool for the detection of subclinical cartilage injury, which would facilitate early therapeutic intervention prior to the development of irreversible degenerative joint disease.

Several mechanisms may contribute to the release of extracellular mtDNA from chondrocytes. While cell death is known to result in passive mtDNA release *(16, 54)*, our *in vitro* data suggests that active release of mtDNA in the absence of cell death also occurs after exposure to an inflammatory stimulus. While the mechanisms underlying this active release in cartilage have not yet been investigated, in other tissues, mDAMP release is closely associated with mitochondrial dysfunction *(16, 25)*, which is likely a contributing factor in chondrocytes as well. Prior studies have demonstrated that IL-1β stimulation of chondrocytes causes mitochondrial dysfunction characterized by increased NO production, decreased mitochondrial respiratory chain complex I activity, and membrane depolarization *(9, 55)*. Mitochondrial dysfunction can subsequently increase ROS production within the cell, which has been associated with mtDNA release via activation of the MPTP, alterations in mitophagy, or production of mitochondria-derived vesicles *(24–28)*. Any of these mechanisms may play a role in active release of mtDNA from chondrocytes and further studies would be necessary to identify which processes are most significant in the context of cartilage injury and early joint disease.

Teasing out active versus passive release of mtDNA is complicated by variation in the intracellular mtDNA:nDNA ratio of chondrocytes. Osteoarthritis chondrocytes have been shown to contain decreased mtDNA content, as reflected by a decreased intracellular mtDNA:nDNA ratio, which can be attributed to mitochondrial dysfunction and impaired mitochondrial biogenesis *(56)*. Furthermore, studies in several species have found that mitochondrial content and function decrease with age *(57)*. Variability in intracellular mtDNA content confounds changes detected in the extracellular mtDNA:nDNA ratio when cell death causes release of the entire DNA content of the cell. Therefore, we normalized the extracellular mtDNA:nDNA ratio to the intracellular mtDNA:nDNA ratio for a given population of chondrocytes to better differentiate between active and passive mtDNA release. To the authors’ knowledge, this method of normalization has not been reported previously, but our data suggest it is the most appropriate method for *in vitro* detection of selective mtDNA release from primary cultured cells, as it accounts for changes in intracellular mtDNA content during cell passage, as well as variability between individuals. Beyond *in vitro* studies investigating a single cell type, however, mtDNA:nDNA ratio calculation diminishes in value. Our previous studies demonstrated that chondrocytes experience differential levels of mitochondrial dysfunction in response to an impact injury, with cells near the injury site experiencing higher levels of strain and resultant mitochondrial depolarization than those farther away *(41)*. This spectrum of dysfunction is likely mirrored by a range of intracellular mtDNA turnover (i.e. mitophagy), as well as extracellular release *(58)*. As such, measurement of mtDNA:nDNA ratio would not be expected to accurately reflect changes in mtDNA release at the individual cell level. At the whole joint level, multiple cell and tissue types likely contribute to extracellular mtDNA and nDNA concentrations, making mtDNA:nDNA ratios even more complicated to interpret. The lack of homogeneous cell populations and inability to accurately normalize to intracellular mtDNA;nDNA ratios likely explain the lack of difference between mtDNA:nDNA ratios in our *ex vivo, in vivo*, and clinical samples. Thus, from a practical perspective, measurement of absolute mtDNA concentration likely provides the most useful information.

Importantly, we found that mitoprotective treatment with SS-31 increased spare respiratory capacity of chondrocytes and decreased release of mtDNA from injured cartilage explants. Other groups have evaluated the effects of mechanical stimuli on release of mtDNA. One study found that fibroblasts released mtDNA in response to changes in tissue stiffness, suggesting that mechanotransduction may play an important role in selective mtDNA excretion *(19)*. Mechanotransduction is known to influence chondrocyte function. Specifically, studies have demonstrated strain-dependent mitochondrial dysfunction *(41, 59)* and ROS release *(60)* in chondrocytes. As several of the proposed mechanisms for active mtDNA release occur in response to mitochondrial dysfunction and increased ROS production *(26, 29, 30)*, mechanotransduction may represent an important link between mechanical injury and active mtDNA release. We previously showed in a similar cartilage explant model that SS-31 treatment mitigates peracute mitochondrial dysfunction after impact injury *(41)*. Together, these studies suggest that mitochondrial dysfunction and extracellular mtDNA release occur concurrently in chondrocytes experiencing mechanical stress. Thus, extracellular mtDNA may represent a practical means for detecting sublethal mitochondrial dysfunction in chondrocytes and for evaluating response to mitoprotective therapy following articular injury.

When translating our *ex vivo* findings to an *in vivo* model, we found that increases in synovial fluid mtDNA are detectable after articular injury. Evidence suggests that in addition to its possible utility as a biomarker of mechanical injury, extracellular mtDNA may also act as a signaling molecule that perpetuates joint inflammation in early OA. Due to its bacterial ancestry, mtDNA contains non-methylated CpG islands that can bind and stimulate toll-like receptor 9 in immune cells, triggering a pro-inflammatory signaling cascade *(61–64)*. Collins et al. demonstrated that intra-articular injection of mtDNA produced inflammatory arthritis in mice, whereas injection with nDNA did not *(65)*. Thus, initial release of mtDNA by chondrocytes following mechanical stress may serve to initiate a pro-inflammatory response throughout the joint via activation of immune cells such as synovial macrophages. Interestingly, we found that in our articular injury model, elevations in nDNA were detectable for a longer period of time and showed more pronounced response to mitoprotective therapy than elevations in mtDNA. These findings may support mtDNA as a more specific marker of early joint injury. It is possible that some joints were experiencing low-grade mechanical stress due to poor conformation or athletic activity prior to injury. This low-level stress may have been sufficient to produce mitochondrial dysfunction and increases in baseline (pre-injury) mtDNA concentration but not severe enough to cause increased nDNA concentrations due to cell death. Moreover, the pro-inflammatory potential of mtDNA and, as our data indicate, the large number of copies of mtDNA per cell and in synovial fluid compared to nDNA may make it a more relevant molecule to quantify. Further studies are warranted to investigate the link between extracellular mtDNA and persistent low-grade joint inflammation, which is an important contributing factor to the development of PTOA *(6)*.

Finally, we found that mtDNA concentration is elevated in cases of naturally occurring intra-articular fractures. This is the first study evaluating synovial fluid mtDNA concentration following traumatic articular injury. In a study investigating the association of mtDNA with immune-mediated arthritis, Contis et al. quantified synovial fluid mtDNA from human joints and obtained values ranging from 10^1^ to 10^4^ copies/mL for patients with chronic osteoarthritis compared to values ranging from 10^2^ to 10^7^ copies/mL for patients with rheumatoid arthritis *(21)*. We obtained mtDNA concentrations on the magnitude of 10^5^ to 10^6^ copies/mL in our equine intra-articular fracture patients, which are much higher than those obtained previously for patients with chronic OA. The chronicity, type of joint pathology (intra-articular fracture versus osteoarthritis), species variation (equine versus human), and differences in mtDNA quantification method could contribute to these differences. Moreover, Contis et al. did not compare synovial fluid mtDNA concentrations in OA joints to those in healthy joints.

In our cases of naturally occurring intra-articular injury, we found no correlation between mtDNA concentration and radiographic changes, but a strong association with the degree of cartilage damage detected during arthroscopic evaluation. This finding is significant because racehorses with carpal osteochondral fragments and mild concurrent cartilage damage return to racing at a significantly higher rate than those with more severe cartilage damage *(44, 66)*. However, radiographic and arthroscopic findings are often disparate in early joint disease, with arthroscopic findings tending to be more severe and of more prognostic value *(44, 67–69)*. Thus, synovial fluid mtDNA concentration may provide a less invasive and expensive means to gain valuable prognostic information that could inform therapeutic decisions. Indeed, in our study, synovial fluid mtDNA concentrations in this subpopulation of elite equine athletes (flat-racing Thoroughbreds) was negatively correlated with the number of race starts in the first year after surgery. Given that there is much debate over the most appropriate measure of post-operative performance in racehorses *(70)*, this association should not be over-interpreted; nevertheless, few studies have found associations between biomarkers and performance outcomes, so this result in a small number of cases warrants further investigation to determine the prognostic value of mtDNA in a larger cohort. In addition, we found that patients with fractures of the middle carpal joint had higher synovial fluid mtDNA than those involving the radiocarpal joint, which is interesting because previous studies found that middle carpal joint fractures carry a worse prognosis for return to racing than radiocarpal joint fractures *(71–73)*. Due to the difficulty in obtaining synovial fluid samples from healthy racehorses, we were unable to evaluate if mtDNA concentrations vary between normal middle carpal and radiocarpal joints in this patient population. Therefore, further investigation would be necessary to determine whether our finding of higher mtDNA in middle carpal joint fluid represents a normal variation between joints or reflects subtle differences in the severity of early pathology not detectable using other diagnostic modalities.

Equine athletes represent an ideal model for the study of naturally occurring osteoarthritis, as equine and human joints are similar in terms of cartilage thickness, cell structure, biochemical properties, and mechanical loading *(74, 75)*. Thus, our findings in equine athletes are particularly relevant to the study of PTOA in humans, and analysis of synovial fluid from human patients to determine the association between joint injury, extracellular mtDNA, and OA in people is warranted.

Our data suggest that mitoprotection may represent a valuable new therapeutic strategy to target early pathomechanisms in PTOA. As a mitoprotective agent, SS-31 has been shown to prevent mitochondrial dysfunction and subsequent ROS production and cell death by stabilizing the inner membrane molecule cardiolipin *(14)*. We found that SS-31 treatment prevents extracellular mtDNA release and increases spare respiratory capacity of chondrocytes, which likely reflect its mitoprotective effects. Importantly, SS-31 treatment also significantly reduced cell death in our *in vivo* talar impact model. Mitochondrial dysfunction is a significant contributor to apoptotic and necrotic cell death in the early time course after articular injury and so mitoprotection may be a valuable therapeutic modality to prevent PTOA.

Although this study provides evidence for the utility of mtDNA in the diagnosis and treatment of early joint disease, it has several limitations. First, while our *in vitro* experiments demonstrated that chondrocytes are one cell type capable of releasing mDAMPs following inflammatory stress, our methods did not allow us to identify the cell/tissue source(s) of elevated mtDNA and nDNA in synovial fluid *in vivo*. As such, mtDNA cannot be considered specific to mitochondrial dysfunction in cartilage, and other joint tissues such as synovium, subchondral bone, or infiltrating immune cells likely also contribute to extracellular mtDNA detected in synovial fluid. Moreover, this study did not determine whether mtDNA is merely a marker of joint injury or an active contributor to the low-grade inflammatory response characteristic of OA progression *(6)*. Further studies are warranted to investigate whether the mtDNA/TLR9 signaling pathway facilitates crosstalk between joint tissues and contributes to the inflammatory response following articular injury, as reported in other disease processes *(61–64)*.

In summary, this study represents the first investigation of extracellular mtDNA in association with PTOA. Our findings indicate the mtDNA may be a valuable biomarker of early cartilage injury. Furthermore, our data suggest that mitoprotective drugs are capable of preventing mtDNA release and cell death, representing a promising new therapeutic strategy for the prevention and treatment of PTOA. Measurement of synovial fluid mtDNA concentration could provide a non-invasive method for detecting mitochondrial dysfunction and cartilage damage associated with early PTOA prior to the appearance of radiographic changes.

## Materials and Methods

### Study Design

This study was designed to test the hypotheses that mtDNA is released from chondrocytes following mechanical injury, that mitoprotective treatment reduces mtDNA release, and that mtDNA is elevated in synovial fluid from joints after articular injury. All animal procedures were approved by Cornell University’s Institutional Animal Care and Use Committee (IACUC). *In vitro* experiments were conducted on cultured equine chondrocytes isolated from the femoropatellar joints of healthy young adult horses (2-5 years) or murine chondrocytes isolated from the coxofemoral joint of healthy neonatal mice (5 days). *Ex vivo* experiments were conducted using cartilage explants harvested from the femoral condyles of neonatal bovids. *In vivo* experiments were conducted using a talocrural impact model of PTOA on young adult horses (2-5 years) with normal talocrural joints based on clinical and radiographic examination. For the *in vivo* experiments both joints of one subject in the SS-31 treatment group (IT) were excluded due to markedly abnormal conformation with the potential to alter joint mechanics. One subject in the control group (IN) was euthanized at 5 weeks due to a medical emergency unrelated to the study. Synovial fluid was collected at euthanasia and analyzed with the 6-week samples of the other subjects. All subjects in each respective experiment were randomly assigned to treatment groups. Clinical carpus SF samples were collected unilaterally or bilaterally from uninjured horses (1-21 years) as well as from those with naturally occurring injury of the carpus (2-20 years). Inclusion criteria were determined prior to sample acquisition. For each experiment, the following were considered one experimental unit: one horse for *in vitro* experiments, one explant for *ex vivo* experiments, one joint for *in vivo* experiments and clinical cases. qPCR was performed using three technical replicates which were averaged for subsequent calculations. Arthroscopic scoring was performed under a blinded consensus method by two of the investigators (M.L.D. and L.A.S.). Radiographic scoring was performed by one blinded investigator (M.L.D.). Prospective power analysis was performed for all experiments using preliminary data obtained by our group to estimate effect sizes. Sample sizes were chosen to provide a study power of at least 80%. All outcome measures and statistical models were decided *a priori*.

### In vitro chondrocyte stimulation

Primary chondrocytes previously harvested and banked from normal femoropatellar joints of healthy adult horses (n = 4, 2-5 years old) were cultured under physoxic conditions (37°C, 5% O2, 5% CO2) for two passages. Twelve hours after plating, second passage chondrocytes were rinsed with phosphate buffered saline (PBS) before serum free stimulation media that contained either interleukin-1β (IL-1β, 1ng/mL) or no stressor (NS) was added to each well. Each combination of horse and stimulant was plated in duplicate. Twelve hours after the addition of stimulation media, conditioned media was collected. For one replicate of each condition, the chondrocytes were trypsinized, rinsed with PBS, and centrifuged at 800xg for 5 min. An aliquot of cells from each well (20% of the cells) was removed for cell lysis and intracellular DNA quantification. For the second replicate of each condition, low-glucose chondrocyte media was added, and the cells were incubated for another twelve hours, after which the media and cells were collected in the same manner as the 12-hour time point.

### Ex vivo cartilage impact injury

A bovine explant model of articular injury was used to evaluate mtDNA release following a mechanical stimulus, as previously described *(11)*. Briefly, cartilage explants were obtained from the medial femoral condyles of healthy bovids (n = 3, 1-3 days of age) within 48 hours of euthanasia using an 8 mm biopsy punch. Samples were rinsed with PBS and cultured in cartilage explant media under standard culture conditions (37°C, 20% O2, 5% CO2). Explants were impacted using a spring-loaded impacting device designed to deliver a rapid compression injury (24.0 ± 1.4MPa peak stress; 53.8 ± 5.3 GPa/s peak stress rate) and instrumented with a load cell on the impactor tip to measure impact force. Explants were divided into two hemicylinders. The hemicylinders were then halved again and portions were selected at random to receive SS-31 (elamipretide, 1 uM) treatment at 1 hour or 12 hours after impact. The explants were cultured as above and cartilage conditioned media was collected 1, 3, 5, and 7 days after impact.

### Microrespirometry

Real-time microscale respirometry was used to measure mitochondrial respiratory function and control in cultured chondrocytes. Fourth passage murine chondrocytes previously harvested and banked from the coxofemoral joints of neonatal UBC mCherry mice (Jax stock 017614, n = 7, 5 days old) were cultured in low-glucose murine chondrocyte media under standard culture conditions (37°C, 21% O2, 5% CO2). Chondrocytes were treated with serum-free low-glucose murine chondrocyte either with or without the addition of SS-31 (elamipretide, 1 µM). 24 hours following addition of SS-31, treated and untreated chondrocytes were trypsinized and cryopreserved in chondrocyte freeze media until microrespirometry was performed.

Extracellular acidification rate (ECAR) and oxygen consumption rate (OCR) were measured within each well using an XFe96 Analyzer (Seahorse Biosciences, North Billerica, MA) approximately every 7 minutes for ∼150 minutes total. After baseline respiration was measured 6 times, a mitochondrial stress test was performed using the XF Cell Mito Stress Test Kit (Seahorse Biosciences, North Billerica, MA) according to standard protocols *(39)*; OCR was measured in response to the automated addition of the following i) oligomycin (1.5 µM; 8 measurements), an ATP synthase inhibitor, ii) carbonyl cyanide-4-(trifluoromethoxy) phenylhydrazone (FCCP; 2.0 µM; 5 measurements), a proton circuit uncoupler, and iii) rotenone (0.5 µM) + antimycin A combination (0.5µM; 4 measurements), inhibitors of mitochondrial complexes I and III, respectively. Following mitochondrial stress test, chondrocytes were washed with PBS and stained with calcein acetoxymethyl (4 µM) for 30 min at 37°C in the dark. Each well was fluorescently imaged with an Olympus IX73 inverted microscope, and viable cell count per well was determined using a custom ImageJ macro. Micro-respirometry data was normalized to cell number by dividing oxygen consumption rates in each well by the number of cells per well. Mitochondrial functional indices were calculated as previously described *(39)*: non-mitochondrial respiration (NMR) = rotenone/antimycin A-stimulated OCR; basal OCR (bOCR) = initial OCR – NMR; maximal OCR (mOCR) = FCCP-stimulated OCR – NMR; spare respiratory capacity (SRC) = mOCR – bOCR; proton leak = (oligomycin-stimulated OCR - NMR)/bOCR; ATP production = (initial OCR – oligomycin-stimulated OCR)/bOCR.

### In vivo cartilage impact injury

An equine talocrural model of impact-induced articular injury was used to evaluate mtDNA content in synovial fluid following mechanical injury as previously described *(42)*. Briefly, adult horses (n = 12, 2-5 years) with clinically and radiographically normal talocrural joints were placed under general anesthesia. Three focal cartilage injuries were created arthroscopically on the medial trochlea of both tali using a hand-held impacting device *(76)*. Either the mitoprotective peptide SS-31 (elamipretide, 1 uM, n = 6), or saline (control, n = 6) was administered intra-articularly to both injured joints 1 hour after injury. Synovial fluid was collected from each joint before injury as well as 7, 14, 28, and 42 days after injury.

### Equine naturally occurring intra-articular fracture cases

Clinical cases were selected from equine patients with intra-articular carpal fractures presenting to either Cornell Hospital for Animals in Ithaca, NY, or Cornell Ruffian Equine Specialists in Elmont, NY, and from historic samples already available through the Cornell Veterinary Biobank. Criteria for inclusion included any horse presenting for arthroscopic removal of osteochondral fragmentation of the carpal bones or distal radius. Synovial fluid was obtained from clinical cases as a byproduct of standard clinical care, such as fluid obtained during arthrocentesis or fluid aspirated prior to joint distension for arthroscopy. A standard set of radiographs (5 views) obtained at the time of presentation or by the referring veterinarian were blindly scored in several categories for changes associated with degenerative joint disease (table S3) by one author (M.L.D.). Arthroscopic video footage was blindly consensus scored for lesions associated with the cartilage, subchondral bone, and synovium (table S5) by two authors (L.A.S., M.L.D.). Post-operative performance was evaluated for all racing thoroughbreds. Race records were obtained through Equibase.com and used to determine whether horses returned to racing and to quantify the number of race starts in the first year following carpal arthroscopy. For comparison, synovial fluid was obtained from the radiocarpal or middle carpal joints of healthy adult horses (n = 7) without clinical evidence of joint disease that were euthanized for reasons unrelated to this study.

### DNA quantification

mtDNA and nDNA were quantified by Taqman qPCR on the ViiA 7 real-time PCR system (Applied Biosystems, Foster City, CA, USA) using equine- and bovine-specific primers and probes. Primers were validated *in silico* and *in situ* as per Bustin and Huggett *(77)*. Samples were analyzed in triplicate and the mean of sample triplicates was used in all subsequent calculations. A standard curve was created using isolated amplicons for each gene target and used to calculate DNA concentration (copies/μL) in unknown samples.

### Statistical analysis

All statistical analyses were performed using JMP Pro 15 software (SAS, Cary, NC). For *in vitro* samples, groups were compared using students’ (paired) t test. Intracellular DNA content was compared across horse and passage using a two-way ANOVA. For *ex vivo* experiments, groups were compared across time and treatment using a linear mixed effects model with impact and explant quadrant as random variables. Given the lack of statistical difference between 1-hour and 12-hour treatment times, the two treatment groups were combined for all analyses. Synovial fluid samples from experimentally injured joints were compared across time points using a linear mixed effects model with joint and horse as random variables. For comparison between treated and untreated groups, data were normalized to baseline values by dividing by pre-injury DNA concentrations and compared using a linear mixed effect model with joint as a random variable. For mixed effects models and ANOVAs, Tukey’s post hoc test for multiple comparisons was used. Data from clinical cases of intra-articular fracture were compared using students’ t test or one-way ANOVA. Fisher’s exact test was used to determine associations between categorical variables. Pearson correlation analysis was performed to determine associations between radiographic, arthroscopic, and performance data and DNA content. Analysis of soft tissue swelling/joint effusion scores was performed by stratifying data into scores < 2 (none/mild) and scores >=2 (moderate/severe) and using students’ t test. Of note, this method of stratification was not decided *a priori*. For each model, residuals were evaluated for normality and homogeneity. Data were log transformed as necessary. Two-sided tests were used for all comparisons and significance was set at p < 0.05.

## Supplementary Materials

Materials and Methods

Fig. S1. Intracellular mtDNA:nDNA ratio varies by individual and cell passage.

Fig. S2. mtDNA:nDNA ratio in cartilage conditioned media after impact injury.

Fig. S3. Delaying mitoprotective treatment to 12 hours after injury does not affect nDNA release from cartilage explants.

Fig. S4. mtDNA:nDNA ratio in synovial fluid of intra-articular carpal fracture patients.

Fig. S5. Total arthroscopy score and cartilage lesion depth correlate with synovial fluid nDNA concentration in joints with intra-articular carpal fractures.

Fig. S6. Cartilage lesion depth scoring sites.

Table S1. Intra-articular carpal fracture patient demographics and sample information.

Table S2. mtDNA, nDNA, and mtDNA:nDNA ratio in injured joints by sex, breed, age, and side of patient.

Table S3. Radiographic scoring rubric.

Table S4. Radiographic scoring correlation data.

Table S5. Arthroscopic scoring rubric.

Table S6. Correlation of arthroscopic scoring with mtDNA:nDNA ratio.

Data file S1. Raw data (provided as a separate Excel file).

Movie S1. Arthroscopic footage of talar impact.

## Acknowledgments

The authors thank Matthew Thomas for technical assistance; Heath Manning, Ariel Bohner, Lisa Mitchell, and the Cornell Veterinary Biobank for assistance identifying and obtaining experimental samples; Lynn Johnson and the Cornell Statistical Consulting Unit for statistics consulting; and Dr. Jim Casey, Laura Keller, and Dr. Maureen Bennett for their intellectual input.

## Author contributions

M.L.D. conceived of and initiated the study, L.A.S. and I.G.S. performed *in vitro* experiments and qPCR, K.L.M. performed microrespirometry experiments, and M.L.D performed *ex vivo* and *in vivo* experiments. All authors contributed to data collection, analysis, and interpretation. L.A.S. drafted the manuscript, with contributions from I.G.S and K.L.M., and revision by M.L.D. All authors contributed to editing the manuscript and approve the final submission.

## Data and materials availability

All data associated with this study are present in the paper or the Supplementary Materials.

