## Supplemental Materials for "Synovial Fluid Mitochondrial DNA Concentration Reflects the Degree of Cartilage Damage After Naturally Occurring Articular Injury"

**Supplementary Materials:**

Materials and Methods

*Intracellular DNA quantification for chondrocytes of increasing passage*

Primary chondrocytes previously harvested and banked from normal femoropatellar joints of healthy adult horses (n = 3, 2-5 years old) were cultured in standard chondrocyte media (Ham's F12 containing 10% FBS, HEPES 0.025 mL/mL, penicillin 100 U/mL, streptomycin 100 U/mL, glucose 10mM) under physoxic conditions (37°C, 5% $O_2$, 5% $CO_2$). Chondrocytes were trypsinized when they reached 90% confluency, at which time an aliquot of cells was removed for lysis and intracellular DNA quantification. The remaining cells were replated. This process was repeated through passage 3 and for a total of four replicates per horse.

*Specific culture conditions*

For *in vitro* experiments, chondrocytes were cultured in in low-glucose chondrocyte media (1:1 Ham's F12: DMEM containing 10% FBS, HEPES 0.025 mL/mL, penicillin 100 U/mL, streptomycin 100 U/mL, glucose 5mM) until stimulation. Second passage chondrocytes were plated on a 24-well plate at a density of 0.5 x $10^6$ cells/well and stimulated with serum free stimulation media (1:1 Ham's F12: DMEM containing HEPES 0.025 mL/mL, penicillin 100 U/mL, streptomycin 100 U/mL, glucose 5mM) with or without the addition of interleukin-1β (1ng/mL). For *ex vivo* experiments, cartilage explants were cultured in cartilage explant media (phenol free DMEM containing 1% FBS, HEPES 0.025 mL/mL, penicillin 100 U/mL, streptomycin 100 U/mL, glucose 2.25 mM). For microrespirometry experiments, chondrocytes were cultured in low-glucose murine chondrocyte media (DMEM containing 10% FBS, L-glutamine 2mM, penicillin 50 U/mL, streptomycin 50 U/mL, glucose 5mM). At approximately 80% confluency, media was changed to serum-free low-glucose murine chondrocyte media (DMEM containing 2mM L-glutamine 2mM, penicillin 50 U/mL, streptomycin 50 U/mL, glucose 5mM) with or without the addition of SS-31 (1uM). Chondrocytes were cryopreserved in murine chondrocyte freeze media (80% low-glucose murine chondrocyte media, 10% FBS, 10% DMSO) until microrespirometry was performed. Prior to the respirometry assay, chondrocytes were thawed and plated onto a 96 well microplate (Seahorse Biosciences, North Billerica, MA) at a density of 20,000 cells per well. Chondrocytes were incubated under standard culture conditions (37°C, 21% $O_2$, 5% $CO_2$) in low-glucose murine chondrocyte culture media for 24 hours. After 24 hours, media was changed to assay media (DMEM containing glucose 10mM, pyruvate 1mM, L-glutamine 2mM; Seahorse Biosciences, North Billerica, MA) and the mitochondrial stress test was performed.

*Sample processing and storage*

Following collection, all media samples were centrifuged at 800xg for 5 min to remove cells and stored at -80°C until further analysis. Synovial fluid samples were centrifuged at 1800 xg for 15 min to remove cells and the resulting supernatant was stored at -80°C until further analysis.

*Taqman qPCR*

DNA was prepared using the QIAamp DNA Mini Kit (Qiagen, Germantown, Maryland, USA) according to the manufacturer’s instructions. The following custom primers and probes were designed and validated: equine NADH dehydrogenase subunit 1 (mtDNA): forward 5'-CCCATCATGACCCTTAGCCA-3', reverse 5'-ATGGAGCTCGGTTGGTTTCG-3', probe 5'-AATGTGATTCATCTCAACATTAG-3'; equine glyceraldehyde-3-phosphate dehydrogenase (nDNA): forward 5'-AGGCTCTTTGCTGCCTTTCT-3', reverse 5'-GCAGGATCAGCACCACTTCT-3', probe 5'-CCCTGGGGATCTGGGGAACCAG-3'; bovine NADH dehydrogenase subunit 1 (mtDNA): forward 5'-GCCTACTTCAACCCATCGCC-3'; reverse 5'-GAGGCTGAAGATGTAGCGGG-3'; probe 5'-TCATTAAAGAACCACTACG-3'; bovine glyceraldehyde-3-phosphate dehydrogenase (nDNA): forward 5'-TCGGTAAACAGCCCTTCACACT-3'; reverse 5'-CACCCTGTTGCTGTAGCCGAA-3'; probe 5'-TCCTTCCAGGTACGACAATG-3'. Quantitative PCR was conducted with denaturation at 95°C for 10 min, followed by 40 cycles of 15 s at 95°C and 1 min at 56°C. Samples were analyzed in triplicate and replicates that deviated more than 0.5 $C_T$ from the mean were discarded. The mean of sample triplicates was used in all subsequent calculations. A standard curve was created through amplification of the target sequence for each gene followed by isolation of the amplicon using the QIAquick Gel Extraction kit (Qiagen, Germantown, Maryland, USA) according to the manufacturer's instructions. Amplicon concentration was determined spectrophotometrically (NanoDrop 1000, Thermo Fisher, Wilmington, Delaware, USA) and used to create a standard curve through serial dilution. Unknown samples were compared to the standard curve to determine DNA concentration (CN/µL).

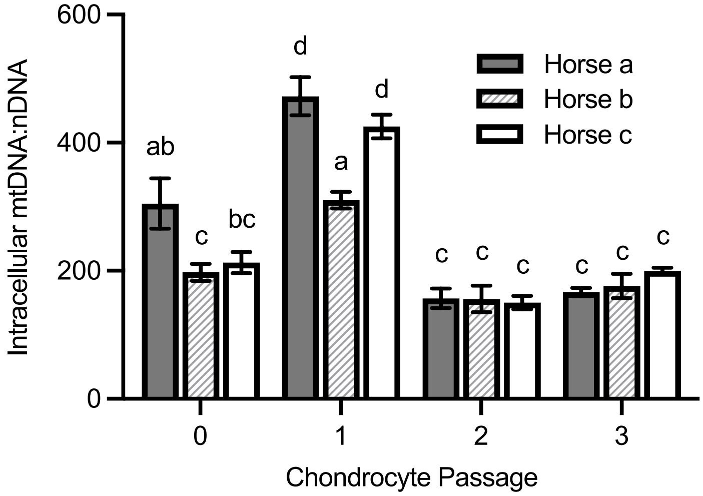


**Fig. S1. Intracellular mtDNA:nDNA ratio varies by individual and cell passage.** Intracellular mtDNA:nDNA ratio measured in primary (passage 0) through third passage chondrocytes of three healthy adult horses. Data are means ± SEM, n = 4, statistics by two-way ANOVA. Groups not sharing a letter are significantly different ($p < 0.05$).

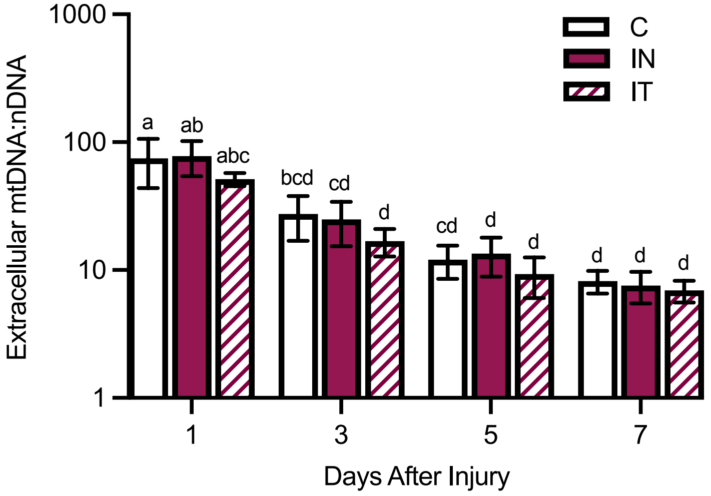


**Fig. S2. mtDNA:nDNA ratio in cartilage explant conditioned media after impact injury.** Cartilage conditioned media mtDNA:nDNA ratio from explants delivered a rapid impact injury and treated with the mitoprotective peptide SS-31 (IT, n = 16) compared to untreated, injured (IN, n = 9) and uninjured, untreated (C, n = 6) explants. Data are means ± SEM, statistics by linear mixed effects model. Groups not sharing a letter are significantly different ($p < 0.05$).

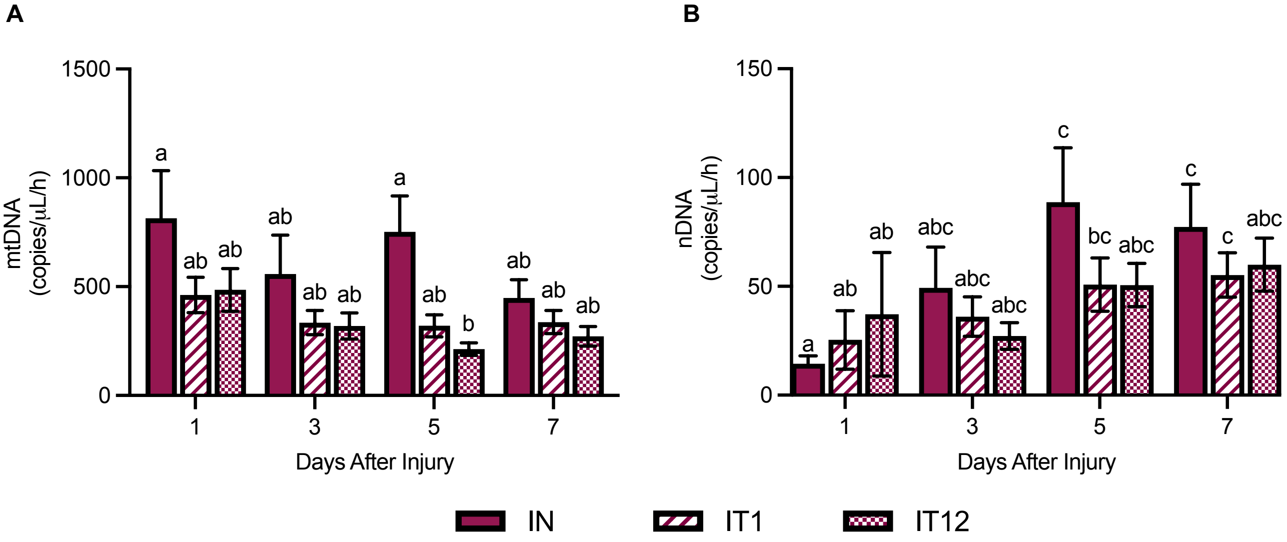


**Fig. S3. Mitoprotective treatment at 12 hours after injury prevents DNA release as effectively as treatment at 1 hour.** (**A-B**) Cartilage conditioned media mtDNA and nDNA concentration from explants treated with the mitoprotective peptide SS-31 at 1 hour (IT1, n = 10) or 12 hours (IT12, n = 6) after rapid impact injury compared to injured, untreated (IN, n = 9) explants. Data are means ± SEM, statistics by linear mixed effects model. Groups not sharing a letter are significantly different ($p < 0.05$).

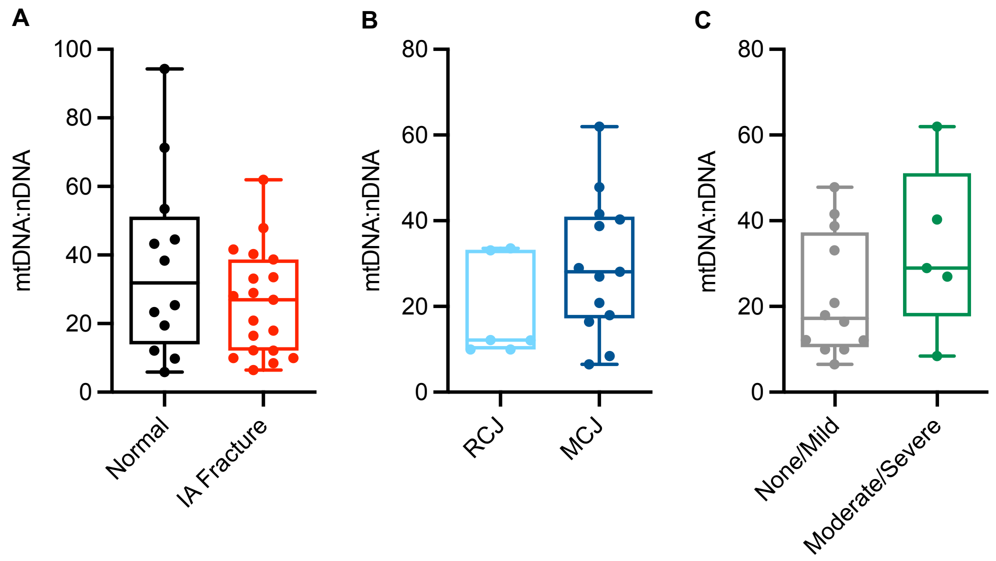


**Fig. S4. mtDNA:nDNA ratio in synovial fluid of equine patients with intra-articular carpal fractures.** (**A**) Synovial fluid mtDNA:nDNA ratio in equine carpal joints with intra-articular (IA) fractures (n = 19) compared to normal equine carpal joints (n = 12). (**B**) Comparison of synovial fluid mtDNA:nDNA ratio from radiocarpal (RCJ, n = 6) and middle carpal (MCJ, n = 13) fractures. (**C**) Comparison of synovial fluid mtDNA:nDNA ratio in joints with no or mild soft tissue swelling (n = 12) to those with moderate/severe soft tissue swelling (n = 5). Statistics by students' t test ($p < 0.05$).

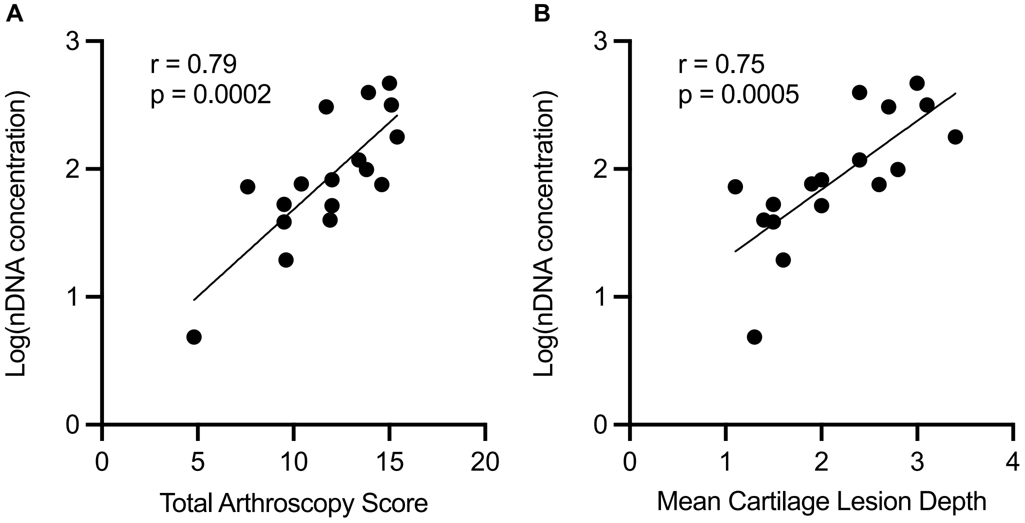


**Fig. S5. Total arthroscopy score and cartilage lesion depth correlate with synovial fluid nDNA concentration in joints with intra-articular carpal fractures.** (**A-B**) Correlation of total arthroscopy score and mean cartilage lesion depth with synovial fluid nDNA concentration in joints with IA carpal fractures. Statistics by Pearson's correlation analysis.

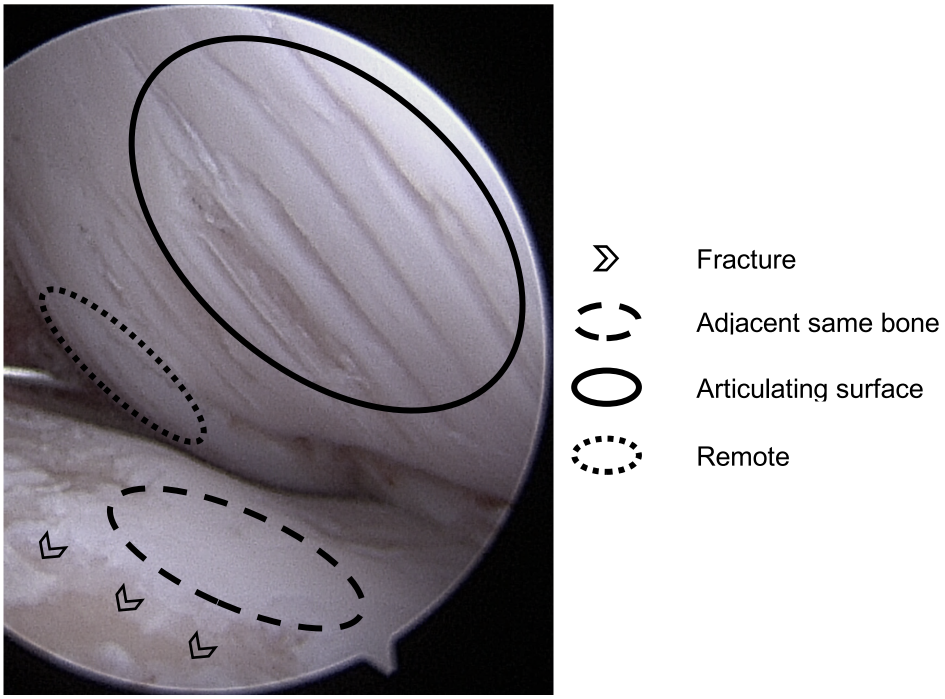


**Fig. S6. Cartilage lesion depth scoring sites.** Cartilage lesion depth was scored at four sites throughout the injured joint: immediately surrounding the fracture, adjacent to and on the same bone as the fracture, on the articulating joint surface to the fracture, and at remote locations within the same joint. These four scores were averaged to obtain a mean cartilage lesion depth score.

**Table S1. Intra-articular carpal fracture patient demographics and sample information.**

NA: information not available, TB: thoroughbred, SB: standardbred, QH: quarter horse, WB: warmblood, RC: radiocarpal joint, MC: middle carpal joint

| Sample | Horse | Group | Sex | Breed | Age (yrs) | Racehorse | Joint | Side |
|---|---|---|---|---|---|---|---|---|
| 1 | A | Fracture | Male, castrated | TB | 4 | Yes | RC | Left |
| 2 | B | Fracture | Male, castrated | TB | 4 | Yes | RC | Left |
| 3 | C | Fracture | Male, intact | TB | 3 | Yes | RC | Left |
| 4 | D | Fracture | Male, castrated | TB | 5 | Yes | RC | Right |
| 5 | E | Fracture | Male, castrated | TB | 4 | Yes | RC | Right |
| 6 | F | Fracture | Male, castrated | TB | 3 | Yes | RC | Right |
| 7 | G | Fracture | Male, castrated | TB | 4 | Yes | MC | Left |
| 8 | H | Fracture | Male, castrated | TB | 4 | Yes | MC | Left |
| 9 | I | Fracture | Male, castrated | TB | 3 | Yes | MC | Left |
| 10 | J | Fracture | Male, castrated | TB | 3 | Yes | MC | Left |
| 11 | K | Fracture | Male, castrated | SB | 5 | Yes | MC | Left |
| 12 | L | Fracture | Male, castrated | QH | 20 | No | MC | Left |
| 13 | M | Fracture | Male, castrated | QH | 4 | No | MC | Left |
| 14 | N | Fracture | Male, intact | TB | 3 | Yes | MC | Right |
| 15 | O | Fracture | Male, intact | TB | 2 | Yes | MC | Right |
| 16 | P | Fracture | Female | TB | 5 | Yes | MC | Right |
| 17 | Q | Fracture | Female | TB | 3 | Yes | MC | Right |
| 18 | R | Fracture | Male, castrated | TB | 6 | No | MC | Right |
| 19 | S | Fracture | Female | SB | 2 | Yes | MC | Right |
| 20 | T | Normal | Female | NA | 15 | No | MC | Left |
| 21 | T | Normal | Female | NA | 15 | No | MC | Right |
| 22 | U | Normal | Female | NA | 8 | No | NA | Left |
| 23 | U | Normal | Female | NA | 8 | No | NA | Right |
| 24 | V | Normal | Male, castrated | NA | 3 | No | NA | Left |
| 25 | V | Normal | Male, castrated | NA | 3 | No | NA | Right |
| 26 | W | Normal | NA | NA | 3 | No | NA | Left |
| 27 | X | Normal | Male, castrated | WB | 21 | No | NA | Left |
| 28 | X | Normal | Male, castrated | WB | 21 | No | NA | Right |
| 29 | Y | Normal | Male, intact | Paint | 1 | No | NA | Left |
| 30 | Y | Normal | Male, intact | Paint | 1 | No | NA | Right |
| 31 | Z | Normal | Female | TB | 1 | No | NA | NA |

**Table S2. mtDNA, nDNA, and mtDNA:nDNA ratio in injured joints by sex, breed, age, and side of patient.** Data are mean ± SD.

| | **mtDNA** (copies/µL) | **nDNA** (copies/µL) | **mtDNA:nDNA** |
|---|---|---|---|
| Sex | | | |
| Stallion (male, intact) | 1763 ± 1414 | 195 ± 240 | 13 ± 7 |
| Gelding (male, castrated) | 3542 ± 4122 | 128 ± 126 | 29 ± 16 |
| Mare (female) | 3182 ± 3510 | 106 ± 70 | 25 ± 13 |
| Breed | | | |
| Thoroughbred | 3463 ± 3980 | 136 ± 128 | 25 ± 13 |
| Standardbred | 2702 ± 2806 | 58 ± 25 | 40 ± 31 |
| Quarter Horse | 1767 ± 2233 | 201 ± 277 | 24 ± 21 |
| Age | | | |
| <3 years | 1873 ± 1634 | 254 ± 303 | 12 ± 8 |
| 3-5 years | 2573 ± 2451 | 89 ± 73 | 28 ± 15 |
| >5 years | 9275 ± 8384 | 357 ± 55 | 28 ± 28 |
| Side | | | |
| Left | 2937 ± 2325 | 120 ± 128 | 31 ± 15 |
| Right | 3501 ± 4875 | 151 ± 148 | 20 ± 14 |

**Table S3. Radiographic scoring rubric.**

| **Criteria** | **0** | **1** | **2** | **3** |
|---|---|---|---|---|
| **Periarticular osteophytes** | None | Mild (1, <2mm) | Moderate (1-2, 2-4mm) | Severe (>2, >4mm) |
| **Subchondral bone sclerosis** | None | Mild | Moderate | Severe |
| **Subchondral bone lysis** | None | Mild (focal shallow lysis) | Moderate (diffuse shallow lysis) | Severe (deep lysis) |
| **Joint space narrowing** | None | Mild (<25% narrowing) | Moderate (25-75% narrowing) | Severe (>75% narrowing) |
| **Fracture** | None | Mild (complete fracture, no comminution, involves one articular surface). Overall impression of acute based on appearance of fragment(s). | Moderate (complete fracture, mild to moderate fragmentation, opposing articular surface mildly affected). | Severe (complete fracture, severe fragmentation, opposing articular surface moderately to severely affected). Overall impression of chronicity based on fragments. |
| **Soft tissue swelling/joint effusion** | None | Mild | Moderate | Severe |

**Table S4. Radiographic scoring correlation data.** Statistics by Pearson's correlation analysis.

| | **mtDNA** log(concentration) | | **nDNA** log(concentration) | | **mtDNA:nDNA** | |
|---|---|---|---|---|---|---|
| | **R** | **P-value** | **R** | **P-value** | **R** | **P-value** |
| Periarticular Osteophytes | -0.36 | 0.151 | -0.16 | 0.528 | -0.37 | 0.144 |
| Subchondral Bone Sclerosis | -0.05 | 0.852 | 0.10 | 0.715 | -0.29 | 0.255 |
| Subchondral Bone Lysis | -0.13 | 0.622 | 0.09 | 0.733 | -0.39 | 0.124 |
| Joint Space Narrowing | -0.32 | 0.209 | -0.16 | 0.533 | -0.34 | 0.186 |
| Fracture Severity | -0.30 | 0.240 | 0.01 | 0.971 | -0.61 | 0.008 |
| Total | -0.27 | 0.293 | -0.02 | 0.954 | -0.49 | 0.046 |

**Table S5. Arthroscopic scoring rubric.**

| **Criteria** | **0** | **1** | **2** | **3** | **4** |
|---|---|---|---|---|---|
| **Lesion depth** | No significant abnormalities | Nearly normal (soft indentations, mild fibrillation and/or superficial fissures) | Mild-moderately abnormal (lesion extending down to <50% cartilage depth) | Moderate-severely abnormal (lesions extending down to >50% cartilage depth but not through subchondral bone) | Severely abnormal (lesions extending down through subchondral bone) |
| **Lesion area** | No significant abnormalities | Focal lesion, <5% of articular cartilage examined | Lesions comprise 5-25% of articular cartilage | Lesions comprise 25-50% of articular cartilage | Lesions comprise >50% of articular cartilage |
| **Subchondral bone** | No significant abnormalities | Abnormal (avascular, yellow-tan discolored, brittle) SC bone closely associated with fracture fragment only, minimal bone loss | Focal area (<10% articular surface) abnormal SC bone not directly associated with fragment, mild bone loss | Multifocal or regional area (10-30% of articular surface) of abnormal SC bone, moderate bone loss | Widespread (>30% of articular surface) and severely abnormal SC bone, extensive bone loss |
| **Active synovitis** | No significant abnormalities. | Mildly abnormal; few hyperemic fronds (<5%, adjacent to fracture site). | Mild-moderately abnormal; <25% hyperemic fronds. | Moderately abnormal; 25-75% hyperemic fronds. | Severely abnormal: > 75% visible fronds hyperemic. |
| **Chronic synovial change** | No significant abnormalities | Mildly abnormal; rare (<5%) thickened/fibrotic fronds. | Mild-moderately abnormal; few (<25%) thickened/fibrotic fronds. Mildly increased volume. | Moderately abnormal; 25-75% visible fronds thick and fibrotic. Moderately increased volume. | Severely abnormal; >75% visible fronds thick and fibrotic. Severely increased volume and/or nodular proliferation. |

**Table S6. Correlation of arthroscopic scoring with mtDNA:nDNA ratio.** Statistics by Pearson's correlation analysis.

| | mtDNA:nDNA | |
|---|---|---|
| | **R** | **P** |
| Cartilage Lesion Depth | 0.27 | 0.303 |
| Cartilage Lesion Size | 0.05 | 0.862 |
| Subchondral Bone | 0.18 | 0.484 |
| Active Synovitis | 0.19 | 0.467 |
| Chronic Synovial Change | -0.16 | 0.545 |
| Total | 0.14 | 0.580 |
